# VCFCons: a versatile VCF-based consensus sequence generator for small genomes

**DOI:** 10.1101/2021.02.26.433111

**Authors:** Elizabeth Tseng, Qiandong Zeng, Lax Iyer

**Affiliations:** Pacific Biosciences, Menlo Park, CA, USA; Laboratory Corporation of America Holdings, Westborough, MA 01581, USA

## Abstract

We had developed VCFCons to address urgent need for a robust consensus sequence generator for SARS-CoV-2 viral surveillance, which presented several unique requirements, including: (a) low coverage areas should be noted with ‘N’s, (b) low frequency or suspicious variant calls need to be filtered. We have found that, while some existing tools such as bcftools can generate the desired consensus sequence, it required multiple filtering steps and additional scripting. VCFCons can generate consensus sequences based on variant calls in a VCF format with versatile filtering criteria based on coverage and estimated variant frequency. We applied VCFCons to the Labcorp SARS-CoV-2 sequencing data and showed that it generated correct consensus sequences that were successfully submitted to GISAID and NCBI. We hope the community will find value in this tool and aim to continue developing VCFCons to handle more complex viral data in the future.

## Introduction

Generating consensus sequences plays an important role in sequencing data analysis. Consensus sequences are generally created with reference-based approaches. The first approach uses a VCF file (Danecek, et al., 2011) as the input to build the consensus sequences by comparison with the reference, as is done by bcftools (Li, 2011) The second approach, used by CLCBio, iVar (Grubaugh, et al., 2019) and Racon (Vaser, et al., 2017) use read alignment (BAM) files or read pileup as the input and create the consensuses directly. A third approach, employed by the pbaa (PacBio amplicon analysis, https://github.com/PacificBiosciences/pbAA) tool, uses a reference-guided (but not reference-aligned) clustering approach to generate consensus sequence, which can then be converted into a VCF file to populate additional quality information for variant filtering.

In our efforts to generate consensus sequences for SARS-CoV-2 from sequencing data reflecting the variant calls, we found existing tools to be inadequate to address our needs. Though many of the tools produced satisfactory consensus sequences, we found the following issues with the consensus sequence generation process. Racon does not accept a VCF as an input or a threshold below which an ‘N’ base should be used to indicate inadequately covered bases. CLCBio consensus generated an incorrect number of ‘N’s, resulting in frameshift errors in the consensus sequence. iVar can only generate consensus sequences from an aligned BAM file, not a VCF. bcftools can accept a VCF file but ignores allele depth (AD) information and requires a separate mask file to produce ‘N’s in low-coverage regions.

As a result, we have developed VCFCons to generate consensus sequences from a VCF file with an accompanying BAM read depth file. VCFCons uses VCF variant calls as the input, but also takes into account the base-level reference coverage info, plus variant call quality and ALT allele frequency.

## Methods

VCFCons generates a consensus sequence based on the following criteria:

- Variant call: variant calls are provided in VCF format. Currently acceptable calls include substitutions, insertions, and deletions.
- Coverage: for positions with a called variant, coverage is determined by the VCF DP column; for positions without a called variant, coverage is provided through an accompanying BAM read depth file (such as those generated by samtools depth). User may determine the minimum coverage threshold below which a ‘N’ instead of the REF (if no variant at this position) or ALT (if variant at this position) is generated.
- QUAL: user may determine the minimum QUAL threshold below which the variant is ignored.
- ALT frequency: the frequency of the variant (ALT) is provided in the VCF file using the DP and AD columns. User may determine the minimum ALT frequency below which the variant is ignored.

VCFCons has been tested with VCF files produced by the CLC variant caller, bcftools, DeepVariant (Poplin, et al., 2018), and pbaa (PacBio amplicon analysis). As shown in Fig 1a, different variant callers produce variable VCF information.

**Fig 1.**
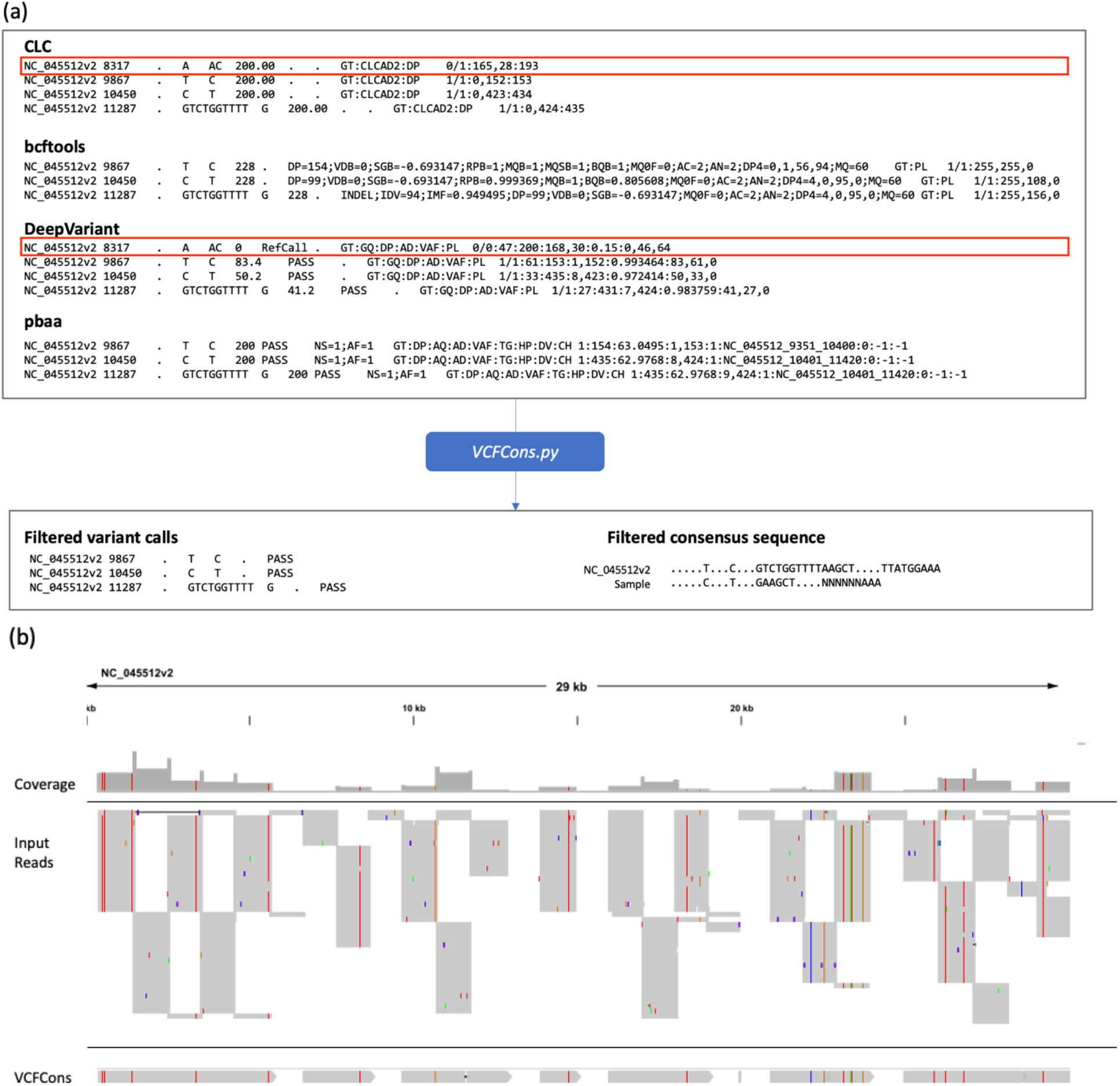
(a) Filtering different variant callers VCF output for SARS-CoV-2 data. The position 8317 has a low frequency (15%) read support, and is not called by bcftools, DeepVariant, or pbaa. All three other calls are jointly supported by all four callers. VCFCons will filter the 8317 call in CLC output based on the variant frequency. VCFCons generates a filtered output VCF file and a consensus sequence. (b) VCFCons consensus sequence visualized in IGV, where low coverage regions are filled with ‘N’ and will not show up in the alignment.

### Code Availability

VCFCons is available at https://github.com/Magdoll/CoSA/

## Results

We compare the results of VCFCons with bcftools and iVar. We did not compare against Racon because it cannot generate ‘N’s, nor can it accept VCF as input. Given the same VCF input file (which is the filtered VCF produced by VCFCons), bcftools (‘bcftools consensus’) and VCFCons both generate the correct consensus sequence. However, bcftools requires an additional step of creating filtered VCFs since it does not read in AD/DP information and also needs masked regions information to produce ‘N’ bases. iVar (‘ivar consensus’) can only accept a pileup file, not a VCF, and further produces an incorrect consensus sequence if there are large indel variants. For example, one of the Labcorp SARS-CoV-2 samples contained a 320bp deletion and the final consensus sequence should be 29583bp based on the Wuhan reference. Both bcftools and VCFCons generated 29583bp consensus sequence, while iVar generated 29903bp, since the pileup is unable to properly detect the large deletion. In summary, VCFCons requires fewer steps to generate the same consensus sequence as bcftools, though both generate accurate results, and can handle large indels and produce ‘N’ bases based on a variety of VCF formats, while other tools fail to produce correct consensus sequences.

## Conclusion

We’ve developed a Python script that can generate consensus sequences for SARS-CoV-2 data based on variant calls in a VCF format with versatile filtering criteria based on coverage and estimated variant frequency. Currently, VCFCons assumes each VCF output contains one major species and will only output one consensus sequence. Future updates will take into account minor species in viral sequencing data as well as potential phased variant information.

## Acknowledgements

Brian Krueger for review and inputs to improve the manuscript and Labcorp and CDC COVID sequencing teams for providing the genomes which are also available in GISAID and NCBI

## Software and References

samtools https://github.com/samtools/samtools

bcftools https://github.com/PacificBiosciences/pbAA

DeepVariant https://github.com/google/deepvariant

pbaa https://github.com/PacificBiosciences/pbAA

Racon https://github.com/isovic/racon

iVar https://andersen-lab.github.io/ivar/html/manualpage.html

QIAGEN CLC Genomics Server and Workbench https://digitalinsights.qiagen.com

## Notes

### Competing Interest Statement

Elizabeth Tseng is an employee of Pacific Biosciences. Qiangdon Zeng and Lax Iyer are employees of Labcorp.

